# MicroRNA-155 regulates osteogenesis and bone mass phenotype via targeting S1PR1 gene

**DOI:** 10.1101/2022.02.18.480982

**Authors:** Zhichao Zheng, Lihong Wu, Zhicong Li, Ruoshu Tang, Hongtao Li, Yinyin Huang, Zhitong Ye, Dong Xiao, Xiaolin Lin, Gang Wu, Richard T Jaspers, Janak L. Pathak

## Abstract

MicroRNA-155 (miR155) is overexpressed in various inflammatory diseases and cancer, in which bone resorption and osteolysis are frequently observed. However, the role of miR155 on osteogenesis and bone mass phenotype is still unknown. Here, we report a low bone mass phenotype in the long bone of miR155-Tg mice compared with control mice. In contrast, miR155-KO mice showed a high bone mass phenotype. miR155-KO mice showed robust bone regeneration in the ectopic and orthotopic model, but miR155-Tg mice showed compromised bone regeneration compared with the control mice. Similarly, the osteogenic differentiation potential of bone marrow stromal stem cells (BMSCs) from miR155-KO mice was robust and miR155-Tg was compromised compared with that of control mice. Moreover, miR155 knockdown in BMSCs from control mice showed higher osteogenic differentiation potential, supporting the results from miR155-KO mice. TargetScan analysis predicted S1PR1 as a target gene of miR155, which was further confirmed by luciferase assay and miR155 knockdown. S1PR1 overexpression in BMSCs robustly promoted osteogenic differentiation without affecting cell viability and proliferation. Thus, miR155 showed a catabolic effect on osteogenesis and bone mass phenotype via interaction with the S1PR1 gene, suggesting inhibition of miR155 as a potential strategy for bone regeneration and bone defect healing.

## Introduction

MicroRNAs (miRNAs) are a class of endogenous non-coding RNAs with 18-22 nucleotides length that bind to the 3′-untranslated region of the target gene and regulate the target gene expression (1). miRNAs regulate cell functions such as growth, differentiation, and energy metabolism by silencing the target gene via degradation or translational repression (1, 2). Moreover, miRNAs are also involved in the pathophysiology of various inflammatory diseases and cancers (3-5). Certain miRNAs had been reported to regulate osteogenesis and bone homeostasis (5, 6). MicroRNA-155 (miR155) is one of the best conserved and multifunctional miRNAs that regulate several biological processes and diseases such as tumorigenesis, cardiovascular disease, kidney diseases, etc (7-10). miR155 is upregulated in inflammatory diseases and cancers, including periodontitis, lung cancer, liver cancer, and breast cancer (11-14). Systemic bone loss is frequently observed in patients with inflammatory diseases and cancers (15, 16). However, the role of miR155 on osteogenesis and bone homeostasis is still unclear.

Induced osteoclasts formation/activity and compromised osteogenic differentiation disrupt bone homeostasis causing bone loss (17, 18). Osteoclasts formation and activity are induced during inflammation and cancer (19, 20). miR155 had been reported to induce osteoclastogenesis (21). miR155 knockout (miR155-KO) mice exhibit reduced local bone destruction in arthritis attributed to reduced generation of osteoclasts (22). Osteogenic differentiation of precursor cells results in bone formation and is the key anabolic event of bone homeostasis. Reduced osteogenic differentiation of precursor cells causes low bone mass phenotype increasing the risk of fracture. Osteogenesis is also a key biological process of bone tissue engineering. Various miRNAs targeted approaches had been developed to promote bone regeneration and bone defect healing during bone tissue engineering (23). The role of miR155 on osteogenic differentiation and bone regeneration is rarely investigated. Compromised osteogenesis and low bone mass phenotype are frequently observed in patients with inflammatory diseases and cancers (24, 25). Similarly, effective bone regeneration and bone defect healing are also key challenges in patients with inflammatory diseases. miR155 targets multiple genes to regulate the pathophysiology of a specific disease in a cell-type-specific manner (26). Sphingosine 1-phosphate receptor-1 (S1PR1) is one of the target genes of miR155 (27), which has been reported to positively regulate osteogenic differentiation of precursor cells (28, 29). Therefore, it is wise to explore the involvement of S1PR1 in miR155-mediated effect on osteogenesis.

In this study, we aimed to analyze the effect of different levels of miR155 on osteogenesis and bone mass phenotype using miR155 transgenic (miR155-Tg) and miR155-KO mice. This study also investigated the role of miR155 target gene S1PR1 on osteogenic differentiation of BMSCs. We found a catabolic effect of miR155 on osteogenesis and bone mass phenotype via targeting the S1PR1 gene.

## Results

### miR155-Tg mice showed a low bone mass phenotype

Micro-CT images showed fewer and thinner trabeculae in miR155-Tg mice femur compared to control mice (Fig. 1A). Trabecular bone parameter BV/TV, BMD, Tb.N, and Tb.Th were significantly reduced in miR155-Tg mice compared to control mice (Fig. 1B-1E). In miR155-Tg mice Tb.Sp was significantly increased compared to control mice (Fig. 1F). These results indicate the low bone mass phenotype in miR155-Tg mice.

**Figure 1.**
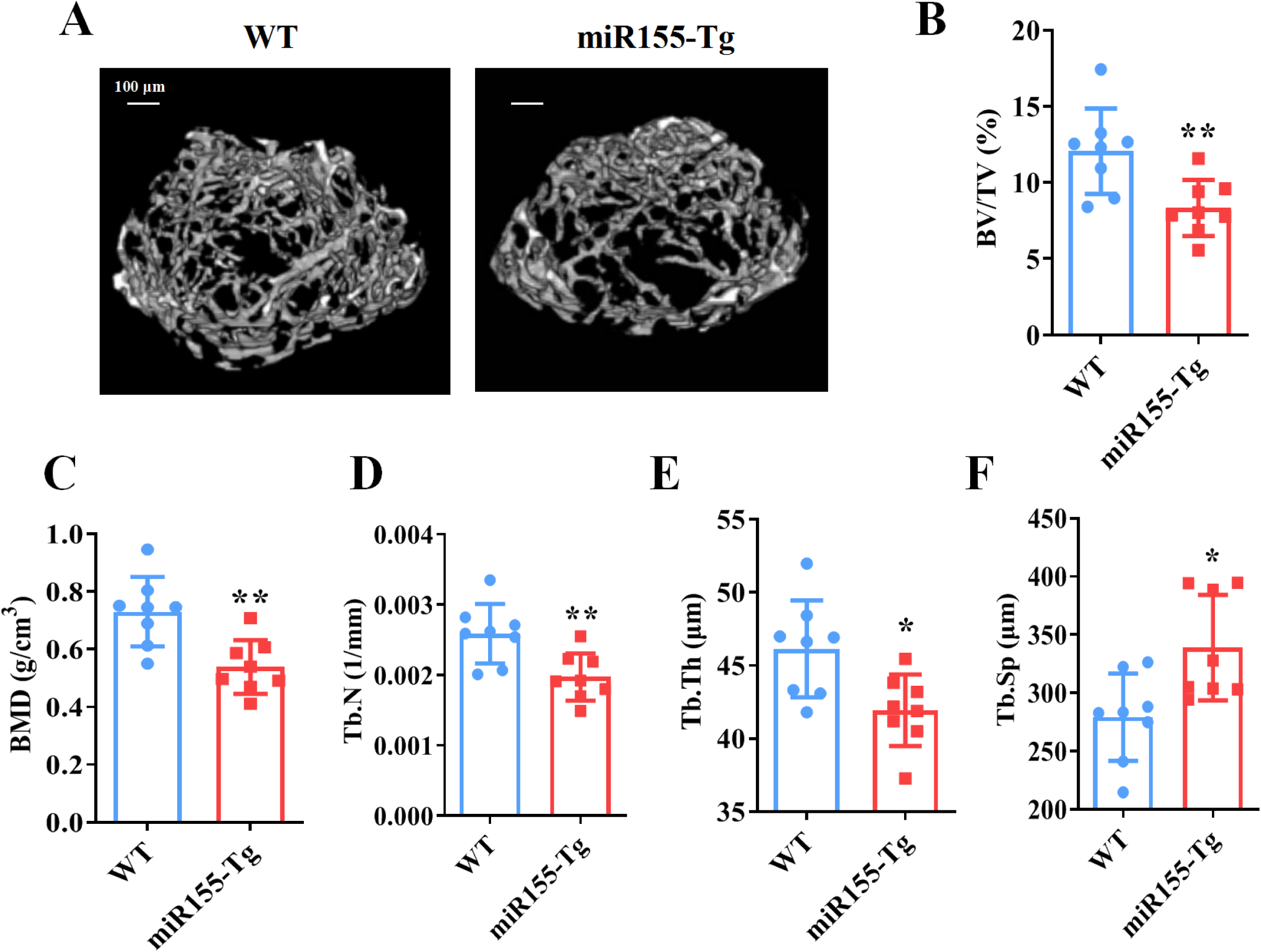
miR155-Tg mice showed a low bone mass phenotype. (A) Representative micro-CT images. (B) BV/TV, (C) BMD, (D) Tb.N, (E) Tb.Th, and (F) Tb.Sp analysis. Data are presented as mean±SD,n=8. Significant difference compared to control mice, *p<0.05, **p<0.01.

### miR155-KO mice showed a high bone mass phenotype

Micro-CT images showed robustly dense and interconnected trabeculae in miR155-KO mice compared to control mice (Fig. 2A). The trabecular bone parameters BV/TV, BMD, Tb.N in miR155-KO mice were 2-, 2.69-, 1.83-, respectively, compared to control mice (Fig. 2B-2D). While Tb.Th was not significantly changed (Fig. 2E). Tb.Sp in miR155-KO mice was significantly reduced compared to control mice (Fig. 2F). These results indicate the high bone mass phenotype of miR155-KO mice. miR155-KO and miR155-Tg showed an opposite trend of bone mass phenotype and trabecular bone parameters (Fig. 1 and 2), suggesting the role of miR155 in bone homeostasis regulation.

**Figure 2.**
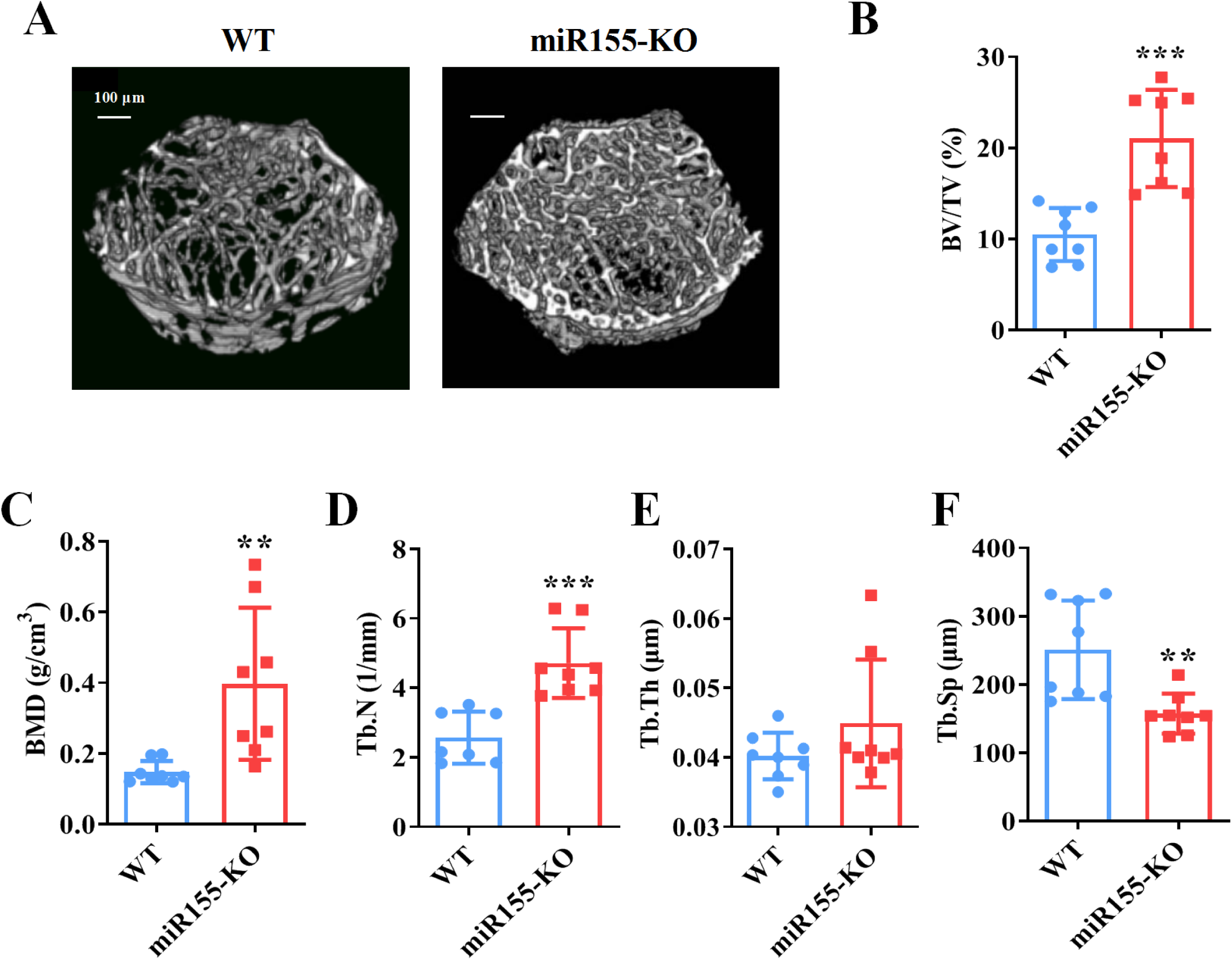
miR155-KO mice showed a high bone mass phenotype. (A) Representative micro-CT images. (B) BV/TV, (C) BMD, (D) Tb.N, (E) Tb.Th, and (F) Tb.Sp analysis. Data are presented as mean±SD,n=8. Significant difference compared to control group, *p<0.05, **p<0.01, ***p<0.001.

### Ectopic bone regeneration was inhibited in miR155-Tg mice while increased in miR155-KO mice

A bone regeneration study was conducted to investigate whether the bone regeneration potential is altered in miR155-Tg and miR155-KO mice. BMP2-loaded collagen membranes were implanted in mice ectopically to confirm the bone regeneration potential in the ectopic site of miR155-Tg, miR155-KO, and control mice. Micro-CT images showed very less bone volume in collagen membrane transplanted in miR155-Tg mice compared to that of control mice (Fig. 3A). The miR155-Tg group showed significantly reduced BV/TV and Tb.N in newly formed bone compared to the control group (Fig. 3B). Tb.Sp was increased in the miR155-Tg group compared to the control group (Fig. 3B). In contrast, ectopic bone regeneration was significantly increased in the miR155-KO group compared to the control group (Fig. 3C). Newly formed bone BV/TV and Tb.N in the miR155-KO group were increased by 6.12-, and 5.64-fold respectively compared to the control group (Fig. 3D). The Tb.Sp was significantly reduced in the miR155-KO group compared control group (Fig. 3J). These results indicate a catabolic effect of miR155 on bone regeneration.

**Figure 3.**
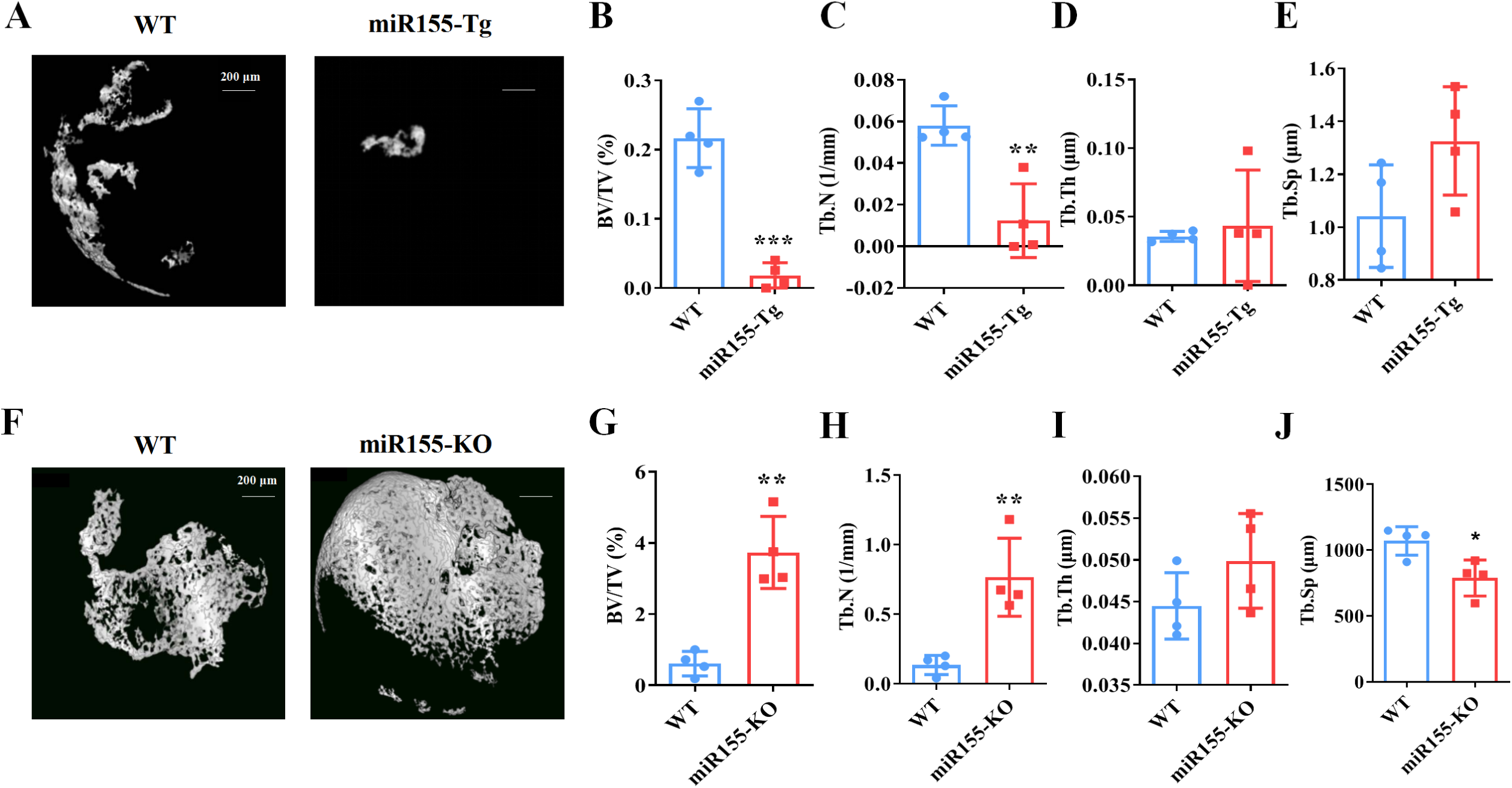
Ectopic bone regeneration was inhibited in miR155-Tg mice but enhanced in miR155-KO mice. (A) Representative micro-CT images. (B) BV/TV, (C) Tb.N, (D) Tb.Th, and (E) Tb.Sp analysis. Data are presented as mean±SD,n=4. Significant difference compared to control group, *p<0.05, ***p<0.001.

### miR155-KO mice showed enhanced bone regeneration in an orthotopic model

To further confirm the catabolic effect of miR155 on bone regeneration, we adopted a calvarial bone defect healing model in miR155-KO mice. Micro-CT images showed more newly formed bone in the defect area of the miR155-KO group compared to the control group (Fig. 4A). Newly formed bone parameter analysis showed higher BV/TV (3.49-fold), and Tb.N (2.85-fold) in the miR155-KO group compared to the control group (Fig. 4B and 4C). There was no significant difference in Tb.Th and Tb.Sp between the groups (Fig. 4D, 4E). These results indicate the anabolic effect of miR155 knockout on bone regeneration.

**Figure 4.**
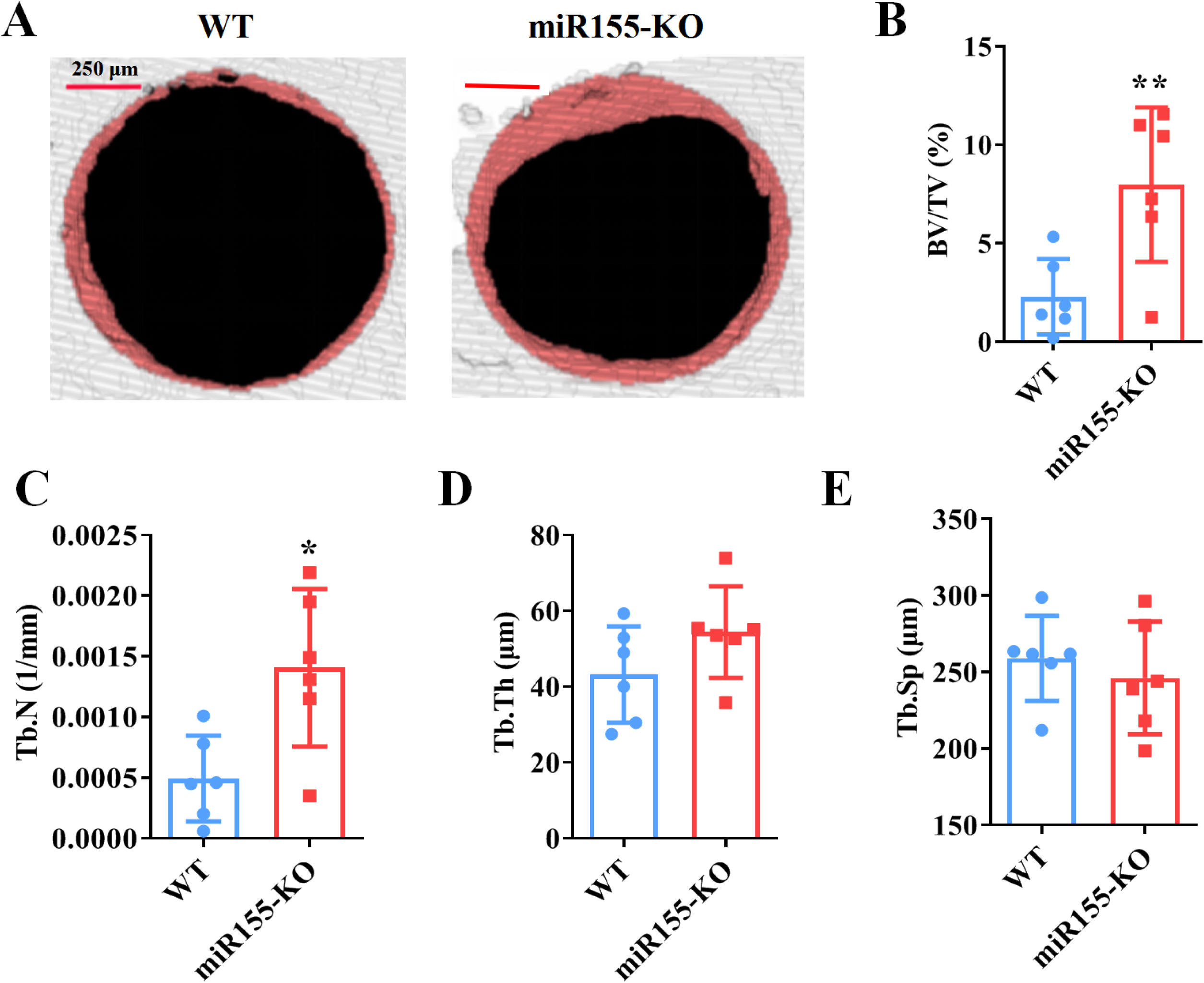
A higher degree of bone regeneration was observed in the calvarial defect of miR155-KO mice. (A) Representative micro-CT images. (B) BV/TV, (C) Tb.N, (D) Tb.Th, and (E) Tb.Sp analysis. Data are presented as mean±SD,n=6. Significant difference compared to control mice, *p<0.05, **p<0.01.

### miR155 influences the osteogenic differentiation of BMSCs

To further confirm the regulatory role of miR155 on osteogenesis, we analyzed the osteogenic differentiation potential BMSCs isolated from miR155-Tg, miR155-KO, and control mice. Mineralized matrix deposition potential in BMSCs from miR155-TG mice was prominently reduced compared to control mice (Fig. 5A and 5B). Similarly, protein level expressions of osteogenic markers ALP and Runx2 in BMSCs from R155-Tg mice were reduced compared to that of control mice (Fig. 5C). These results indicate the compromised osteogenic differentiation potential of BMSCs from miR155-Tg mice. In contrast, BMSCs from miR155-KO mice showed robustly higher matrix mineralization potential compared to that of control mice (Fig. 5D and 5E). The protein level expressions of osteogenic markers ALP were enhanced in BMSCs from miR155-KO (Fig. 5F). These results demonstrated the catabolic effect of miR155 in osteogenic differentiation.

**Figure 5.**
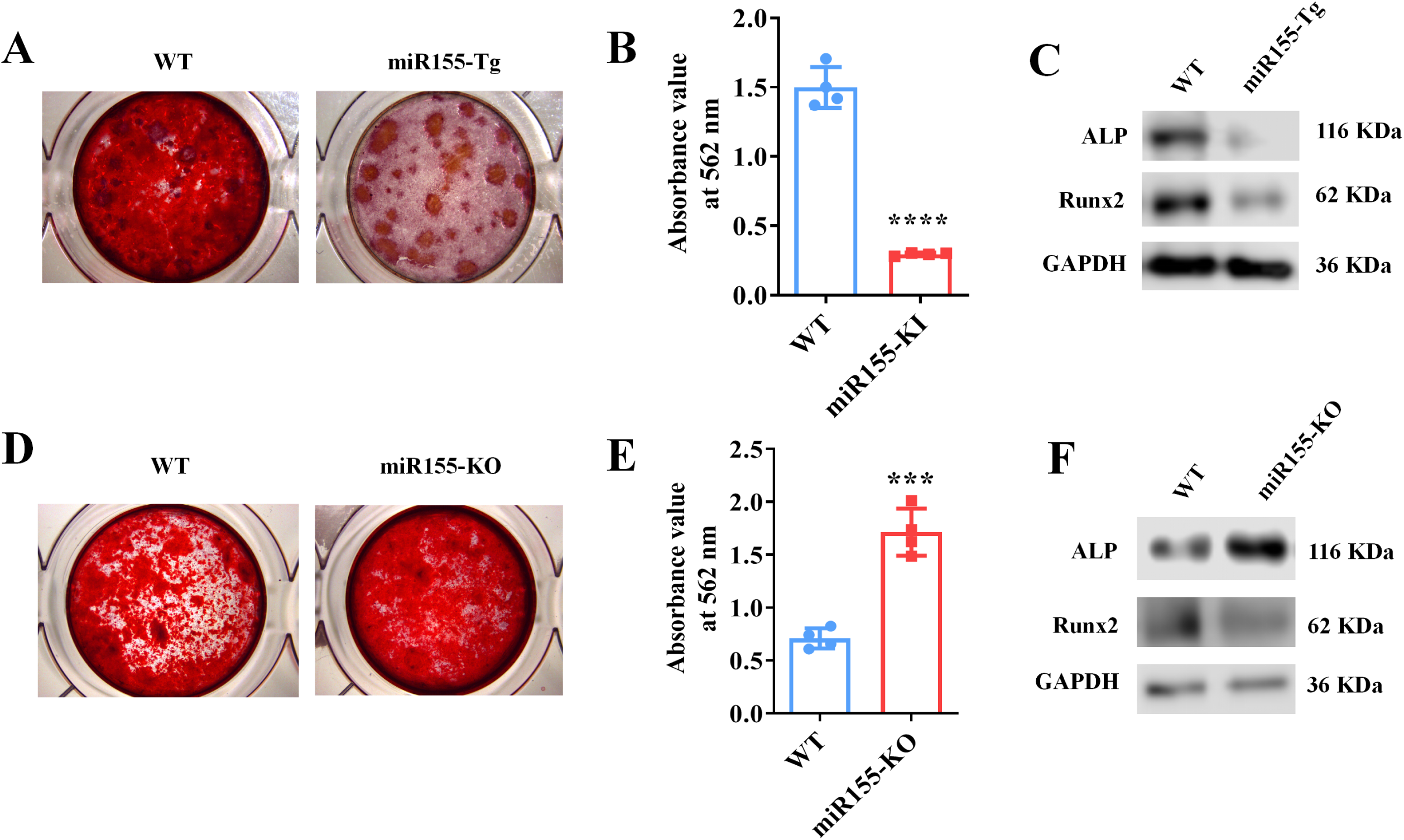
BMSCs from miR155-Tg mice exhibited compromised osteogenic differentiation. (A) ARS images stained at day 10 of culture, (B) ARS quantification, and (C) Western blot analysis of osteogenic markers. BMSCs from miR155-KO mice showed robust osteogenic differentiation potential. (D) ARS images stained at day 10 of culture, (E) ARS quantification, and (F) Western blot analysis of osteogenic markers. Data are presented as mean±SD, n=4. Significant difference compared to control mice, ***p<0.001, ****p<0.0001.

### Knockdown of miR155 induced osteogenic differentiation of BMSCs

Matrix mineralization was robustly increased in the miR155 sponge group compared to the negative control group (Fig. 6A and 6B). miR155 was knockdown in BMSCs to further confirm the role of miR155 level in osteogenic differentiation. miR155 sponge lentivirus treatment significantly reduced the expression of miR155 in BMSCs (Fig. 6C), indicating the successful knockdown of miR155. Similarly, the protein level expression of osteogenic markers ALP and Runx2 was robustly upregulated in the miR155 sponge group compared to the negative control group (Fig. 6E). These results showed a catabolic effect of miR155 on osteogenic differentiation of precursor cells.

**Figure 6.**
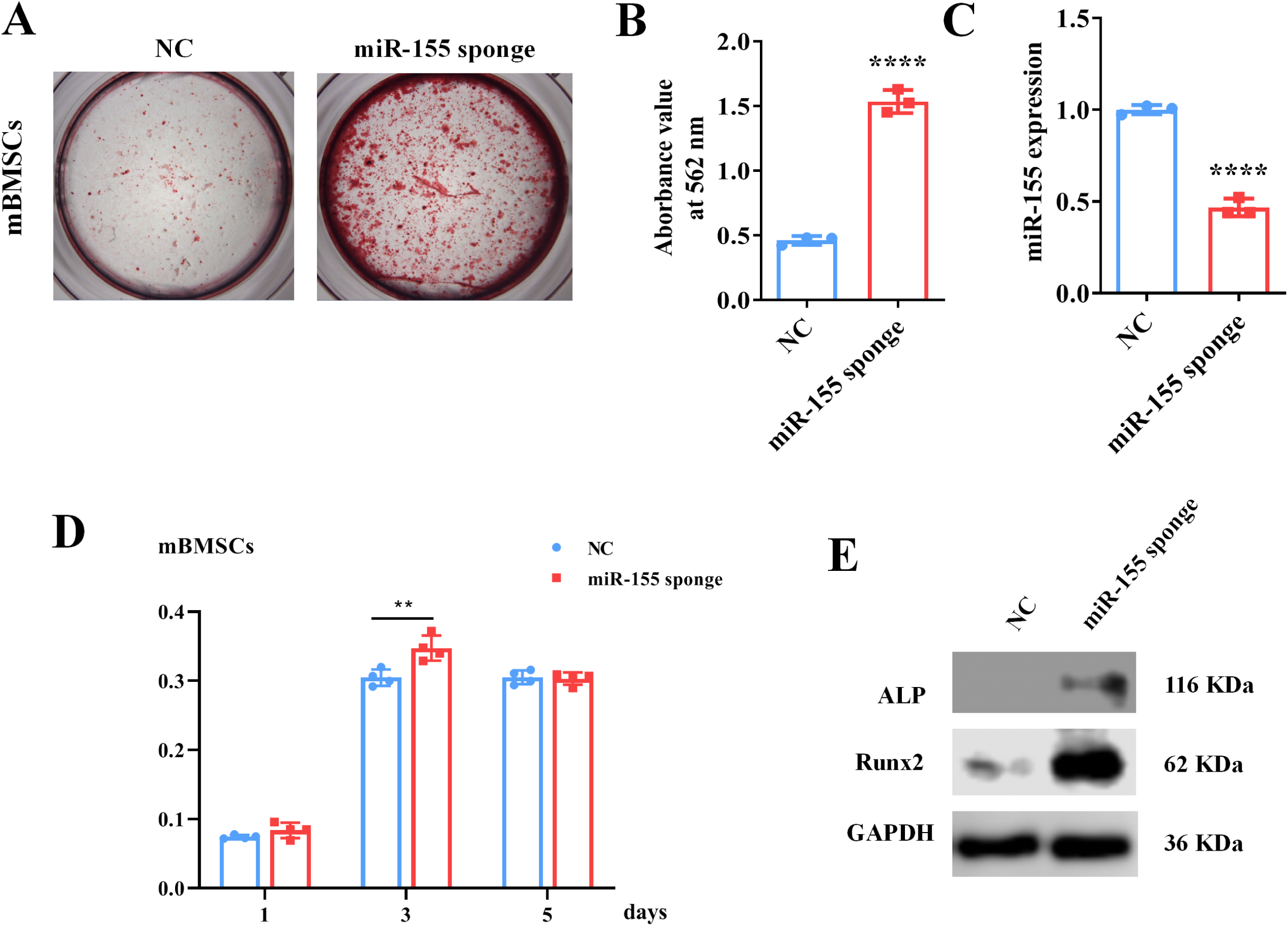
miR155 silenced BMSCs showed higher osteogenic differentiation potential. (A) ARS images stained at 21 days of culture, (B) ARS quantification, (C) miR155 expression level, (D) cell viability, and (E) Western blot analysis. Data are presented as mean±SD. Significant difference compared to negative control, ***p<0.001, ****p<0.0001. NC: negative control.

### miR155 targets the S1PR1 gene to regulate osteogenic differentiation of BMSCs

TargetScan predicted S1PR1 as a target gene of miR155 (Fig. 7A). Luciferase reporter gene assay was performed to analyze the miR155 and S1PR1 gene interaction (Fig. 7B). Our results showed that miR155 directly binds to the 3’UTR of the S1PR1 (Fig. 7B). miR155 robustly enhanced the protein level expression of the S1PR1 gene in BMSCs (Fig. 7C), confirming the interaction of miR155 and the S1PR1 gene. S1PR1 transfection in BMSCs robustly enhanced the matrix mineralization (Fig. 7D and 7E). The protein level expression of S1PR1 was enhanced in lentivirus-mediated S1PR1 overexpressed BMSCs (Fig, 7F). This result indicates the efficacy of lentivirus-based S1PR1 overexpression in BMSCs. The protein level expression of ALP and Runx2 was increased in S1PR1 overexpressed BMSCs (Fig. 7F). S1PR1 overexpression in BMSCs by lentivirus did not affect cell viability and proliferation (Fig. 7G). These results indicate that the miR155 targets the S1PR1 gene to regulate osteogenic differentiation of BMSCs.

**Figure 7.**
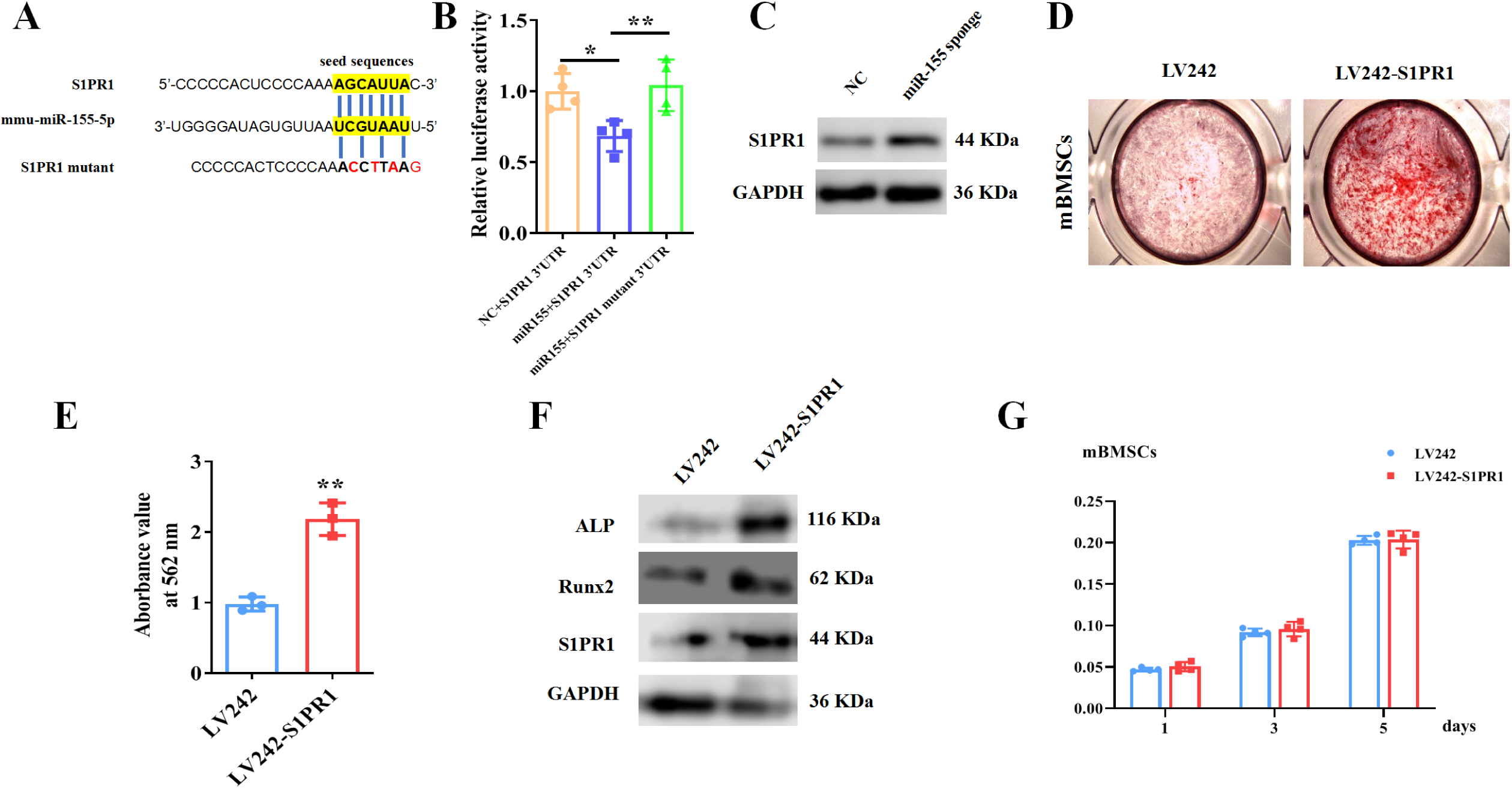
miR155 targets S1PR1 to regulate osteogenic differentiation of BMSCs. (A) The TargetScan prediction of miR155 binding site in S1PR1 gene. (B) Luciferase assay, (C) S1PR1 protein expression in BMSCs, (D) ARS images stained at 21 days of culture, (E) ARS quantification, (F) Western blot analysis of osteogenic markers, and (G) cell viability. Data are presented as mean±SD. Significant difference compared to negative control, *p<0.05, **p<0.001.

## Discussion

Differentiation of mesenchymal stem cells to osteoblasts is a vital event of bone regeneration. Differentiated osteoblasts deposit mineralized matrix and contribute to new bone formation. Various miRNAs had been reported to regulate osteogenesis and bone mass phenotype (23). In this study, miR155-Tg mice showed compromised bone regeneration and low bone mass phenotype. In contrast, miR155-KO mice showed improved bone regeneration and a higher bone mass phenotype. BMSCs from miR155-Tg and miR155-KO mice showed compromised and robust osteogenic differentiation potential, respectively. miR155 silencing also promoted osteogenic differentiation potential in BMSCs. These results indicate a catabolic effect of miR155 on bone regeneration and bone mass phenotype. Knockdown of miR155 in BMSCs robustly enhanced the protein level expression of S1PR1 and osteogenic regulator Runx2, indicating S1PR1 as a target gene of miR155 in BMSCs to regulate osteogenic differentiation. S1PR1 overexpression in BMSCs enhanced Runx2 expression and osteogenic differentiation of BMSCs indicating the regulatory role of miR155-S1PR1 interaction on osteogenesis (Fig. 8).

**Figure 8.**
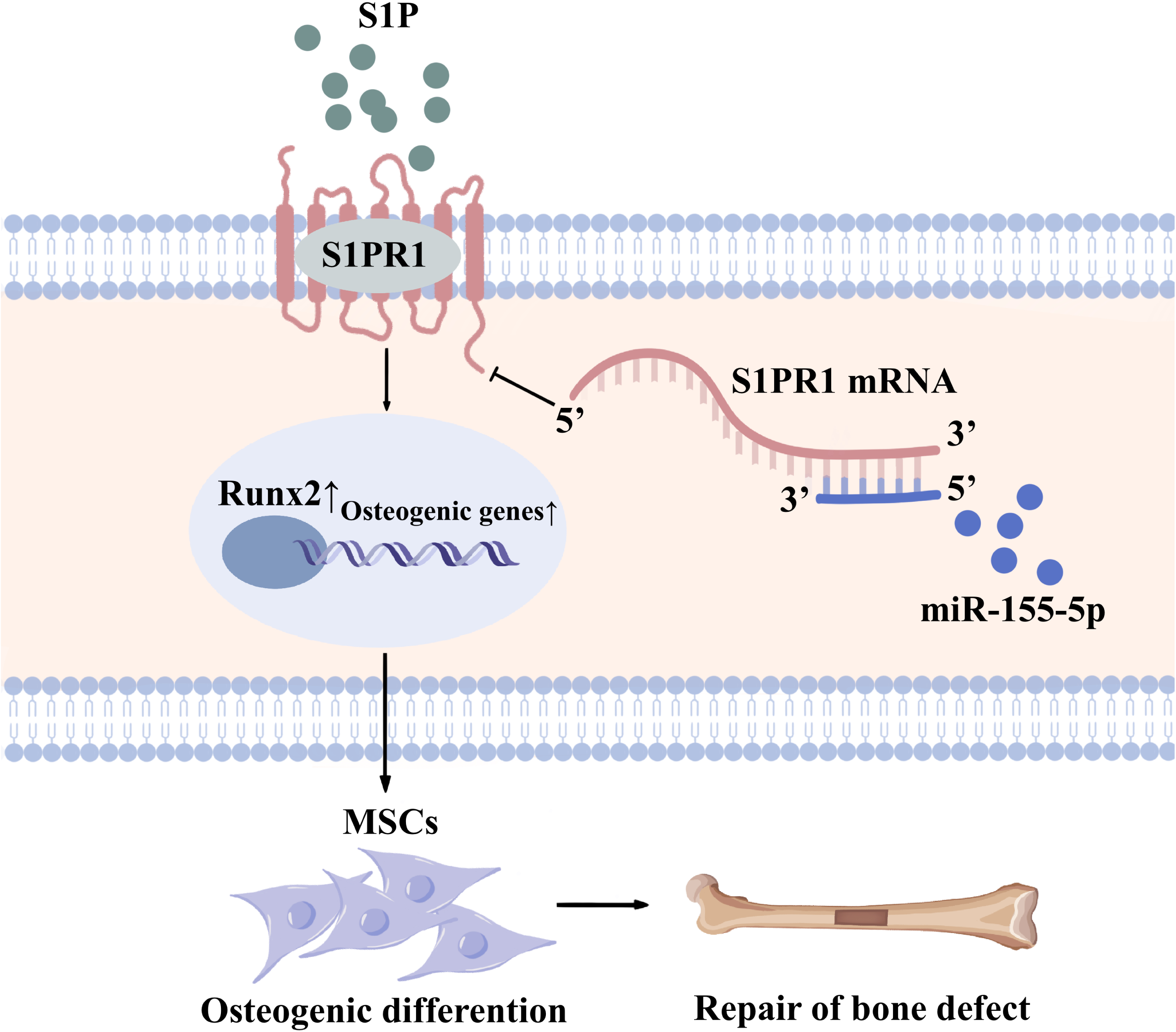
Scheme of miR155-mediated regulation of osteogenesis. S1P activates the S1PR1, further increasing Runx2 expression to regulate the osteogenic differentiation of MSCs into osteoblasts. Finally, activation of S1PR1 signaling increases the bone mass and may be used as repair of bone defects. miR155 inhibits the process by direct binding with 3’UTR S1PR1 mRNAs. MSCs: mesenchymal stromal cells.

miRNAs/anti-miRNAs have been used for bone tissue engineering (23). MiR26a (30), anti-miR31 (31), anti-miR34a (32), miR135 (33), anti-miR138 (34), anti-miR146a (35), miR148a (36), anti-miR221 (37), anti-miR-3555p (38) had shown anabolic effect on osteogenic differentiation of precursor cells and bone defect healing. Most of the miRNAs/anti-miRNAs promote bone regeneration via activation of osteogenic master regulator Runx2 (23). In this study, miR155 overexpression showed a catabolic effect on osteogenesis and bone mass phenotype. Interestingly, knockout or knockdown of miR155 showed an anabolic effect on osteogenesis and bone defect healing. Runx2 was involved in the miR155-mediated regulation of osteogenic differentiation of BMSCs. miR155 had been reported to inhibit osteogenesis of MC3T3-E1 cells via SMAD5 downregulation (39). miR155 inhibits BMP9-induced osteogenic differentiation of precursor cells via downregulation of BMP signaling (40). Qu et al. revealed the miR155 inhibition could alleviate suppression of the osteogenic differnentiation of human BMSCs under the condition of under high glucose and free fatty acids by targeting SIRT1 (41). Our results from miR155-Tg mice on inhibition of osteogenesis are in accordance with the findings from the literature (39, 40). To our knowledge, this is the first study to report the anabolic effect of miR155 knockout or knockdown on osteogenic differentiation of precursor cells. Our results suggest the potential application of anti-miR155 on bone regeneration and bone tissue engineering application.

Anti-miR155 oligonucleotides and antagomir have been designed for various cancer treatments (42, 43). Small molecule-based cyclic peptidomimetics had shown an inhibitory effect on miR155 biogenesis(44). MLN4924 is an inhibitor of the NEDD8-activating enzyme. MLN4924 decreases the binding of NF-κB to the miR155 promoter and downregulates miR155 in AML cells (45). Since, knockdown of miR155 robustly promoted the osteogenesis and bone mass phenotype, anti-miR155 or miR155 inhibitor could apply for bone tissue engineering application as well as the treatment of low bone mass phenotype. However, the osteoinductive potential of already available anti-miR155 or miR155 inhibitors should be tested using in vitro and in vivo models to prove this hypothesis.

miR155 targets different genes in different cells to regulate the cell type-specific functions (46-49). S1PR1, a target gene of miR155, is regulated during various physiological and pathological conditions (49, 50). In this study, miR155 inhibition upregulated S1PR1 protein expression. Overexpression of S1PR1 robustly promoted Runx2 expression and osteogenic differentiation of BMSCs. Gu Y et al. reported that miR155 inhibits osteogenic differentiation of precursor cells via inhibiting SMAD5 (39). Higashi K et al. reported SMAD1/5/8 as downstream signaling of S1PR1/S1PR2 to induce Runx2 expression in osteoblast (29). Reports from the literature and results of this study indicate that miR155 targets the S1PR1 gene to inhibit Runx2 expression thereby reducing bone regeneration and bone mass.

Since miR155 is upregulated in various cancer including hematological cancers (43). Hematological cancer mainly affects bone marrow that is the dwelling of bone precursor cells. Hematological cancers are associated with bone loss and fracture of vertebrae and long bone. Breast and lung cancer are frequently metastasized to bone and cause osteolysis (51, 52). Cancer/cancer metastasis-induced bone loss-mediated fracture is a serious clinical problem. However, the role of upregulated levels of miR155 on cancer/cancer metastasis-related reduced bone mass is still unclear. Moreover, the prevention of cancer/cancer metastasis-induced bone loss is a huge challenge for clinicians. Since anti-miR155 is already proven to be beneficial for cancer treatment and miR155 knockdown promotes bone regeneration, anti-miR155 could treat cancer as well as caner-induced bone loss as a killing two birds with one stone concept. However, future in vitro and in vivo studies are needed to confirm this hypothesis.

miR155 is overexpressed and plays a key role in the pathophysiology of inflammatory diseases including autoimmune arthritis, osteoarthritis, periodontitis (11, 22, 53). An elevated level of miR155 in arthritis promotes M1 macrophage polarization and inflammation (53). Prevention of bone loss in inflammatory diseases using currently available therapeutic approaches is not satisfactory. Moreover, inflammation impedes bone regeneration thereby causing the failure of bone tissue engineering approaches. Since anti-miR155 has anti-inflammatory (54) and bone regenerative potential, anti-miR155 could be potential therapeutic to promote bone regeneration even in inflammatory conditions.

This study used both miR155-Tg and miR155-KO mice to investigate the role of miR155 on osteogenesis and bone mass phenotype. Bone regeneration both in ectopic and orthotopic models confirmed the regulatory role of miR155 in bone regeneration. miR155 silenced and S1PR1 overexpressed BMSCs further confirmed the S1PR1 as a target gene of miR155 to regulate Runx2 expression during osteogenesis. The limitation of this study is that we did not analyze the downstream signaling pathway of S1PR1 that regulates Runx2 expression in BMSCS.

miR155 showed a catabolic effect on osteogenesis and bone mass via targeting S1PR1. Our results suggest miR155 as a potential target to promote bone regeneration and higher bone mass. Since miR155 is overexpressed in inflammatory diseases and anti-miR155 has shown anti-inflammatory potential, the miR155 inhibitors could be potential therapeutics to promote bone regeneration even in inflammatory conditions.

## Materials and methods

### Mice

miR155-KO mice were purchased from the Jackson Laboratory (Stock No. 007745). miR155-Tg mice were constructed as described in our previous reports (55). The C57BL/6J wildtype mice, as the control mice of miR155-KO mice, were purchased from Guangdong medical laboratory animal center. While the FVB mice were littermate control of miR155-Tg mice. The blinded evaluation was used for mice assignments and analysis. The animal experiment was conducted in accordance with the guidelines approved by the Institutional Animal Care and Use Committee of the First Affiliated Hospital of Guangzhou Medical University, Guangzhou, China (2017-078).

### Bone phenotype analysis

Bone phenotype analysis was performed in 8 weeks old male mice (8 mice/group) using micro-CT. Mice were anesthetized using isoflurane (RWD life science CO., China), followed by cervical dislocation. Femur with distal growth plate was collected and fixed in 10% buffered formalin. micro-CT scanning was performed to evaluate bone phenotype using Bruker Sky1172 Skyscan (Kontich, Belgium). A total of 100 slices (1 mm) below the distal growth plate of the femurs was measured for 3D reconstruction and quantification of trabecular bone as described previously (56). The X-ray tube was operated at 96 kV and 65 μA using a 0.5 mm Al filter with a resolution of 7.93 μm pixels. Scanning was performed by 180° rotation around the vertical axis, camera exposure time of 1300 ms, rotation step of 0.6°, frame averaging of 2, and random movement of 10. 3D images were made using CTvox software (Skyscan, Kontich, Belgium). Data viewer software (Skyscan, Kontich, Belgium) was used for images and linear analysis. Relative bone formation parameters including bone volume/total volume (BV/TV), bone mineral density (BMD), trabecular number (Tb.N), trabecular thickness (Tb.Th), and trabecular separation (Tb.Sp) were analyzed.

### Ectopic grafting of collagen membrane

Collagen membrane ZH-BIO (China) with 5 mm diameter and 1 mm thickness were osteogenically functionalized by loading 10 μL of 0.3 mg/mL BMP2 solution. BMP2-loaded collagen membranes were implanted in the subcutaneous pockets of miR155-Tg, miR155-KO, and respective control mice. Eight-week-old male mice with 20-22 g body weight (4 mice/group, 1 membranes/mouse) were used for this study. The subcutaneous transplantation of collagen membrane was performed as described previously (57). After 18 days of transplantation, mice were euthanized by isoflurane, collagen membrane was collected, and further analyzed for newly formed bone using micro-CT.

### Mice calvaria bone defect healing

Eight-week-old male mice with 20-22 g body weight (3 mice/group, 2 defects/mouse) were used for the calvaria bone defect healing study. The calvaria bone defects model was established as previous reports (58, 59). Surgery was performed under anesthesia with pentobarbital sodium (50 mg/kg) and the body temperature was maintained at 37°C with a heated platform. After exposure of the cranium surface through a skin incision, a circular and full-thickness bone defect with a 1 mm diameter was generated across the two sides of the sagittal suture using a trephine drill. The bone fragments were taken out. After gentamicin flushing, the incisions were sewed up. After 1 month, the calvaria bones were collected and the healing level was analyzed by micro-CT.

### The isolation of primary BMSCs

Euthanized transgenic mice and control male mice (5-6 weeks old) were immersed into 75% ethanol for 5 min. The femurs and tibia were acquired. Primary BMSCs were isolated and expanded as described previously (60). In brief, bone marrow was flushed out from the tibia and femurs and disturbed into small pieces. Cells were collected by centrifugation and plated into flasks and allowed to adhere for 24 h. Nonadherent cells were washed, and culture was continued in DMEM supplemented with 10% non-heat inactivated FBS and 1% penicillin/streptomycin. The cells were cultured in a 5% CO_2_ incubator maintaining a humid atmosphere. Cells were trypsinized from 80% confluent culture and passaged. The primary BMSCs in passages 1-3 were used throughout the study.

### Analysis of target gene of miR155

TargetScan software was used to predict the target gene of miR155. TargetScan predicted S1PR1 as a possible target gene of miR155.

### Plasmid construction and lentivirus preparation

The S1PR1 3’UTR sequences and mutant sequences (200 bp upstream and 200 bp downstream of the binding site from NM_007901.5 transcript) were synthesized and cloned into wildtype plasmid pmirGLO Dual-Luciferase miRNA Target Expression Vector (Promega, USA) by Generay (China). We created S1PR1 3’UTR and S1PR1 mutant 3’UTR plasmids for luciferase assay. The LV242 S1PR1 and control LV242 plasmid were purchased from Genecopedia (USA). miR155 negative control (NC) and miR155 sponge plasmids were purchased from OBIO (China).

For lentiviral transduction, HEK293T cells were co-transfected with expression plasmids (miR155 NC, miR155 sponge, LV242, or LV242 S1PR1) with the packaging plasmids pMD2.VSVG, pMDLg/pRRE, and pRSV-REV using EZ trans transfection regent (Shanghai life iLab BioTechnology Co., LTD, China). After 48 h, fresh lentiviral supernatant was collected and used for infection. BMSCs cells were expanded to 60% confluence prior to lentiviral infection. After infection for 10 h, cells were washed and allowed to recover for 24 h and used for subsequent experiments. miR155 sponge efficacy was analyzed by RT-qPCR. S1PR1 overexpression efficacy was analyzed by western blot analysis.

### Luciferase assay

Luciferase assay was performed for further confirmation of S1PR1 as a target gene of miR155. S1PR1 3’UTR (100 ng) with NC (50 nM), S1PR1 3’UTR (100 ng) with miR155 (50 nM), and the S1PR1 mutant 3’UTR plasmid (100 ng) with miR155 (50 nM) were co-transfected into HEK293T cells by lipofectamine 2000 (Thermo Fisher Scientific Inc. USA). After 48 h, the luciferase assay was performed according to the manufacture’s instruction using Luc-Pair™ Duo-Luciferase Assay Kit 2.0 (Genecopedia, USA). In brief, the cells were lysed and incubated with 100 μL Fluc work solution for 5 min, the fluorometric measurement was performed by Varioskan^®^ Flash (Thermo Fisher, USA). The lysis solution was added with 100 μL Rluc, incubated for 5 min, and the fluorometric value was measured.

### Alizarin red staining

BMSCs (28,000 cells/well) were seeded at 48 well culture plate and cultured with osteogenic medium (50 μg/mL vitamin C, 0.01 μM dexamethasone, and 10 mM β-glycerophosphate). Then the cells were fixed with paraformaldehyde and stained with Alizarin Red S solution (1%, pH 4.2) (Solarbio life sciences, China) for 10 min. The staining was visualized under stereomicroscope Leica EZ4HD (Leica, Germany). For quantitative analysis, the alizarin red-stained mineralized matrix was dissolved with 200 μL 10% hexadecylpyridinium chloride monohydrate for 1 h, and the supernatant was collected. The optical density of the supernatant (100 μL) was measured by a microplate reader at 562 nm.

### Immunoblotting

Cells were lysed by using RIPA Buffer (CWBio, China) containing protease inhibitor to extract total protein. Total protein (20 μg) was added to 10% SDS-polyacrylamide gel. The protein was transferred to PVDF membranes (Millipore, USA) after electrophoresis and blocked for 1 h with blocking buffer (Beyotime, China). Then PVDF membranes were incubated with the primary antibodies including ALP (CST, USA, 1:3,000), Runx2 (CST, USA, 1:2,000), S1PR1 (Abcam, UK, 1:2,000), and GAPDH (CST, USA, 1:5,000) overnight at 4°C. The membranes were further incubated with horseradish peroxidase-conjugated secondary antibody for 1 h and reacted with ECL (Millipore, USA). Finally, the photographs were taken by the C-Digit system (Agilent, USA).

### RT-qPCR

The microRNAs were extracted from BMSCs with the MolPure^®^ Cell/Tissue miRNA Kit (Yeasen, China) as the manufacture’s instructions. The microRNA was further reversed by Tailing reaction using miRNA 1st strand cDNA synthesis kit (Accurate biology, China). In brief, 3.75 μL microRNA, 5 μL 2× miRNA RT Reaction Solution, 1.25 μL miRNA RT Enzyme Mix were incubated at 37°C for 1 h, 85°C for 5 min. RT-qPCR was performed using SYBR^®^ Green Premix Pro Taq HS qPCR Kit (Accurate biology, China) on an AriaMx Real-time quantitative PCR machine (Agilent, USA). The PCR reaction conditions were 95°C for 30 s, followed by 40 cycles at 95°C for 5 s and 60°C for 30 s. The fold change relative to the control group was measured by the 2^-ΔΔCt^ method. The primer used for miR155 detection was TAATGCTAATTGTGATAGGGGT.

### Prestoblue cell viability assay

Cell viability was analyzed using PrestoBlue™ cell viability reagent (Thermo Fisher, USA). BMSCs (4×10^3^ cells/well) were seeded into 96 well culture plates. After 1, 3, and 5 days, the medium was removed and replaced with a cell viability detection medium according to the manufacturer’s instructions. After 2 h, the OD value was measured by a microplate reader at 570 nm with a reference wavelength of 600 nm.

### Statistical analysis

Data are expressed as mean±SD. Statistical analysis was performed with t-tests for comparison of two groups. p<0.05 was considered as a significant difference.

## Acknowledgments

This project was funded by National Natural Science Foundation of China (82150410451), the General Guiding Project of Guangzhou (20201A011105), the Medical Scientific Research Foundation of Guangdong Province (B2020027), the Undergraduate Science and Technology Innovation Project of Guangzhou Medical University (2020A049), and High-level University Construction Funding of Guangzhou Medical University (02-412-B205002-1003017 and 06-410-2106035).

## Data availability

Source data files have been provided for Figures 1-7 as Figure 1 source data-1, Figure 2 source data-2, Figure 3 source data-3, Figure 4 source data-4, Figure 5 source data-5, Figure 6 source data-6, Figure 7 source data-7. And the the western blot raw data was provided as source data-8.

